# Optogenetic stimulation of Purkinje cells in the cerebellar vermis disrupts innate freezing behaviors and is highly aversive

**DOI:** 10.64898/2026.04.22.720220

**Authors:** Raven A. McGann, Christopher E. Vaaga

## Abstract

The generation of adaptive defensive behaviors in response to predator threats requires the integration of sensory inputs by neural circuits to shape context-appropriate motor outputs. While the cerebellum is increasingly recognized as a contributor to non-motor behaviors, its role in regulating innate fear behaviors is only beginning to be recognized. For example, it is unclear whether and how cerebellar activity influences both the expression and experience-dependent habituation of defensive responses to ethologically relevant stimuli. Here, we examine how manipulation of cerebellar output alters innate freezing behavior and its adaptation across repeated predator-like visual stimuli. Using optogenetic stimulation of vermal Purkinje cells, we demonstrate that ongoing fastigial nucleus (FN) activity is required for the generation of appropriate freezing behaviors and further that perturbations of FN activity alter adaptive habituation across repeated trials at both short (5 minute) and long (24 hour) intervals. Our results suggest that cerebellar stimulation results in an elevated fear state, as stimulation was both anxiogenic in an open field arena and resulted in robust real-time place aversion that resisted reversal learning. Together, these findings identify the cerebellum as a key regulator of both the expression and experience-dependent adaptation of innate fear responses.

## Introduction

The cerebellum has long been studied in the context of motor control and coordination (Ito, 1984; Apps & Garwicz, 2005); however, numerous lines of evidence suggest that the cerebellum plays an additional important role in non-motor function, including cognition, affective regulation, and language processing (Sacchetti *et al*., 2005; Strick *et al*., 2009; Zhu *et al*., 2011; Apps & Strata, 2015; Adamaszek *et al*., 2017). In fact, in humans, damage to the cerebellar vermis results in a constellation of cognitive and affective deficits which have collectively been referred to as the Cerebellar Cognitive Affective Syndrome (CCAS; (Schmahmann, 2004, 2021; Hoche *et al*., 2018; Ahmadian *et al*., 2019). These observations motivate defining how cerebellar circuits engage in both motor and cognitive/affective regulation. In the motor domain, sensorimotor errors result in climbing fiber recruitment, driving real-time correction of ongoing movements as well as engaging long-lasting synaptic plasticity in the cerebellar cortex, necessary for updating motor programs (Hartell, 2002; D’Angelo, 2018; Hull & Regehr, 2022). The extent to which similar cerebellar computations occur in non-motor domains remains an open question. In human fMRI studies, Crus I/II show elevated BOLD signal in response to unexpected sentence structures, consistent with a role in processing linguistic prediction errors (Lesage *et al*., 2017). However, it is unclear whether and how cerebellar circuits utilize ‘errors’ to guide both ongoing and future fear/affective behaviors.

Although not traditionally viewed as a component of the distributed brain circuits underlying fear behaviors, the cerebellum is extensively interconnected with limbic structures, suggesting the cerebellum plays an important role in fear processing (Apps & Strata, 2015). Fear can be broadly categorized as learned (i.e. conditioned) or innate (i.e. predator threats). In rats, lesions of the cerebellar vermis result in a robust reduction in freezing responses to natural predators, with little impact on conditioned fear (Supple *et al*., 1988; Koutsikou *et al*., 2014), indicating that the cerebellum plays an important role in innate fear regulation. Looming visual stimuli, which mimics an attacking aerial predator, has been used as a model to study innate fear processing in mice (Yilmaz & Meister, 2013; De Franceschi *et al*., 2016; Tafreshiha *et al*., 2021; Qi *et al*., 2025; Carroll *et al*., 2025). Interestingly, in naïve mice, the initial presentation of a looming visual stimulus results in strong/persistent freezing behavior; however, repeated presentations of identical stimuli result in robust habituation of fear behaviors (Tafreshiha *et al*., 2021; Qi *et al*., 2025; Carroll *et al*., 2025). Such habituation may be mediated by a threat-danger ‘mismatch’ (i.e. ‘error’) that engages cerebellar learning and ultimately results in updated fear appraisal.

Generation of motor-related fear behaviors (i.e. freezing or flight) ultimately requires activation of the midbrain periaqueductal gray (PAG;(Bandler *et al*., 1985; Bandler & Carrive, 1988; Zhang *et al*., 1990; Bandler & Shipley, 1994; Walker & Carrive, 2003; Gross & Canteras, 2012). Anatomically, the fastigial (medial) cerebellar nucleus projects to the ventrolateral column of the PAG (vlPAG; (Gonzalo-Ruiz *et al*., 1990; Vaaga *et al*., 2020; Frontera *et al*., 2020). The vlPAG has been shown to preferentially drive freezing behavior and play additional roles in positive prediction errors (Tovote *et al*., 2016; Walker *et al*., 2019; Vaaga *et al*., 2020). While a small proportion of freezing pre-motor neurons receive direct excitatory input from the fastigial nucleus, the predominant effect of fastigial inputs to the vlPAG is modulatory in nature (Vaaga *et al*., 2020). Functionally, fastigial inputs preferentially target a local population of dopaminergic neurons in the vlPAG, which, in turn, modulate synaptic strength in freezing-related premotor neurons. More specifically, either fastigial or dopamine neuron activation increases the strength of synaptic inhibition and decreases the strength of synaptic excitation in freezing-related premotor neurons (Vaaga *et al*., 2020). Together, these data suggest that cerebellar activity may mechanistically contribute to the habituation observed after repeated presentations by altering synaptic integration in freezing related premotor neurons. Such input would therefore be predicted to reduce the efficacy with which sensory stimuli drive fear behavior.

Here, we tested this prediction by optogenetically perturbing activity in the fastigial nucleus during looming visual stimuli. Strikingly, optogenetic stimulation of vermal Purkinje cells at the onset of the looming stimulus significantly impaired the ability of mice to engage in looming-evoked freezing, while also reducing habituation across trials. However, optogenetic stimulation prior to the looming stimulus resulted in robust and consistent freezing responses across trials, suggesting a heightened fear state. Consistent with this observation, alterations in cerebellar output additionally resulted in an anxiogenic phenotype in an open field arena and significant real-time place aversion. Together, these results suggest that ongoing cerebellar activity is necessary for engaging appropriate freezing behaviors and further that disrupting cerebellar activity is, in itself, highly aversive.

## Methods

### IACUC approval and animal use

All animals were housed in accordance with, and all experimental methods were approved by, Colorado State University (protocol 3836/7255, CEV) or Northwestern University (protocol IS00014844) Institutional Animal Care and Use Committees. Animals were obtained from Jackson Laboratories or bred in a satellite housing facility located in the same building as the behavioral testing suite. Adult (>p49) male and female animals on a C57BL/6J genetic background were used for this study. When possible, all experimental cohorts were sex-balanced. Non-transgenic animals served as wildtype controls. All experimental animals (referred to as L7-cre::Ai32) were bred via crossing homozygous Ai32 mice (RCL-ChR2(H134R)/EYFP; Stock #024109) with homozygous L7Cre-2 mice (B6.129-Tg(Pcp2-cre)2Mpin/J; Stock #004146), resulting in the selective expression of ChR2-eYFP in cerebellar Purkinje cells. All mice were socially housed (2-5 mice per cage), when possible, and maintained on a 12:12 hour light:dark cycle with *ad libitum* access to food and water.

### Fiber Optic Cannula Implantation

Unless otherwise noted, all animals (wildtype control and L7-cre::Ai32) underwent stereotaxic surgery to implant a fiberoptic cannula (200 μm core; NA = 0.22) into the midline cerebellar vermis (**Figure 1 A, B**). Mice (>p49) were anesthetized with isoflurane (1-2% to effect) and positioned on a stereotaxic platform. The head was shaved and cleaned with betadine and ethanol. 0.2 mL of 2% lidocaine was subcutaneously injected at the dorsal cranium as a local anesthetic. Following a craniotomy, either a Doric Lenses or Thorlabs 2mm mono fiber-optic cannula was placed within the cortex of the anterior cerebellar vermis using the following coordinates (in mm from bregma); -6.8 posterior, ±0.0 lateral, and -1.4 deep from cerebellar surface. The cannula was secured and the exposed skull fully covered using dental cement. Buprenorphine Extended Release (Bup-ER, 0.6-1.0 mg/kg) was perioperatively administered via a subcutaneous injection for analgesic support. Following anesthetic recovery, mice were closely monitored for at least 72 post-operative hours and allowed at least one week of recovery before onset of behavioral testing.

**Figure 1:**
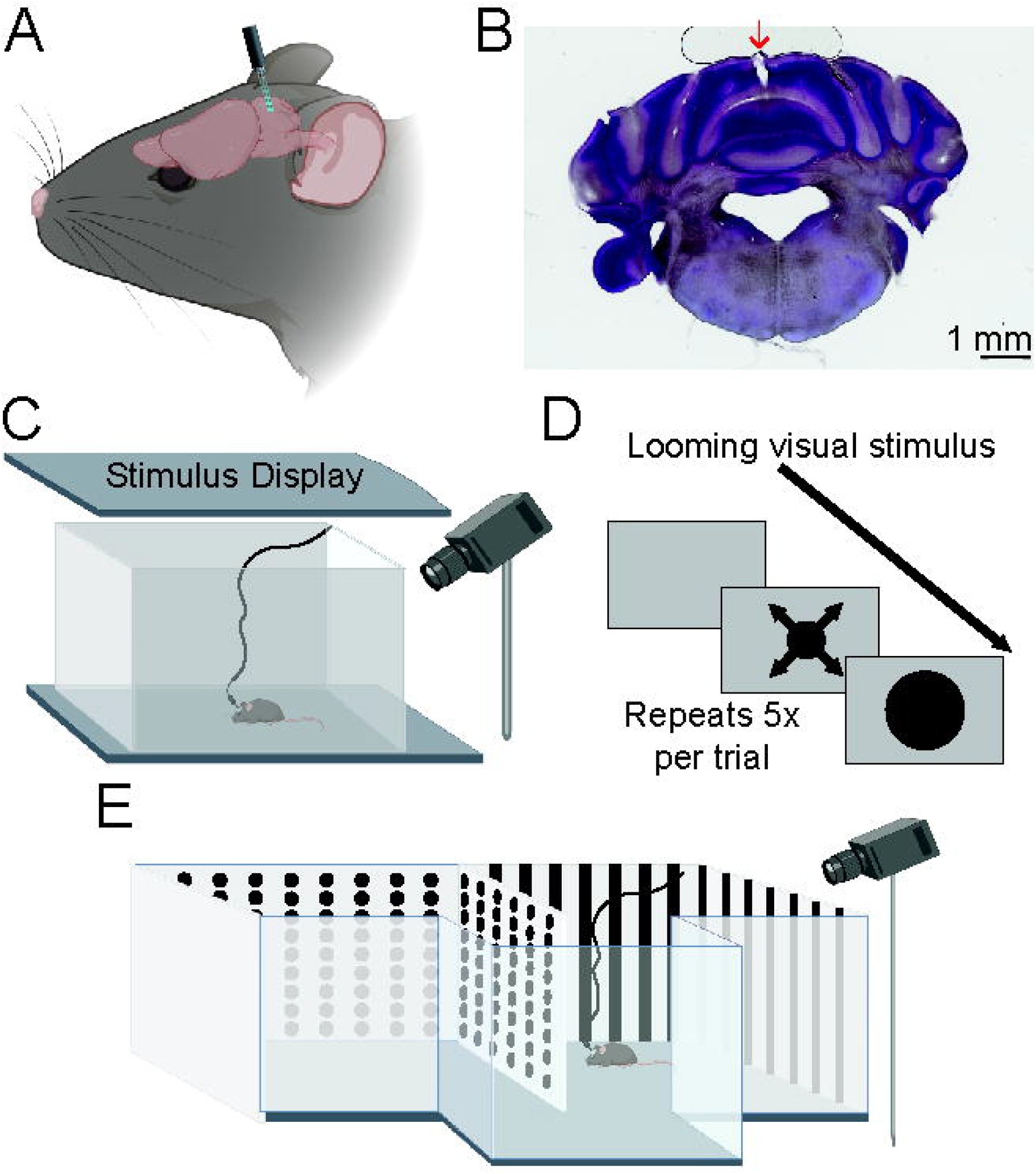
Experimental Methodology. (**A**) Schematic of fiberoptic cannula placement within cerebellar vermis. (**B**) Histological image confirming location of fiber-optic cannula (red arrow) within lobule IV/V of the cerebellar vermis (**C**) Diagram of open-field behavioral arena with overhead monitor used to display the looming visual stimulus. (**D**) Depiction of looming visual stimulus used to evoke innate fear responses. (**E**) Diagram of three-chambered place preference arena consisting of two large, visually distinct rooms interconnected by a clear, plexiglass antechamber.

### In-vivo recordings

To validate the effects of Purkinje cell optogenetic stimulation on fastigial nucleus (FN) firing rates, loose cell-attached recordings were performed in awake, head-fixed mice (Brown & Raman, 2018). Prior to *in vivo* recordings, mice underwent surgical preparation to allow head fixation and repeated access to the cerebellum. Mice were anesthetized with 1– 2% isoflurane (to effect), and lidocaine (2%, subcutaneous) was administered under the scalp for local anesthesia. Buprenorphine ER-Lab (1 mg/kg, subcutaneous) was administered perioperatively to provide extended postoperative analgesia.

For head fixation, two small stainless-steel screws (1/16 SL; Fisher) were inserted into the parietal bones to improve implant stability. A custom headplate was then positioned over the skull and secured using dental cement. A craniotomy was subsequently made over the medial cerebellar nucleus (relative to bregma: −6.25 mm posterior, ±0.6 mm lateral). The exposed brain surface was covered with silicone elastomer (Kwik-Sil, World Precision Instruments) to protect the craniotomy.

Following surgery, mice were allowed to recover for at least one week prior to experimentation. Animals were then habituated to head fixation for 3–4 days before electrophysiological recordings were performed. Loose cell-attached recordings of FN neurons were made in awake head-fixed mice using borosilicate glass pipettes (3-6 MΩ) filled with Tyrode’s solution (in mM: 150 NaCl, 4 KCl, 2 CaCl_2_, 2 MgCl_2_, 10 HEPES, 10 glucose; pH 7.35 with NaOH). Recordings were amplified/digitized using a Multilcamp 700B/Digidata 1550B and acquired using pClamp. Recordings from the FN were made from A-P: -6.2 to -6.5, M-L: 0.6 – 1.3 (in mm from bregma) and at a depth ranging between 1.8 and 3.1 mm. CbN neuron identity was confirmed using three criteria: spontaneous firing, silenced by optogenetic stimulation of Purkinje cells, and the lack of complex spikes (Brown & Raman, 2018; Brown *et al*., 2024). For optogenetic stimulation, a fiber optic cannula (200 μm core, NA = 0.22, ThorLabs) was placed on the surface of the cerebellum at the midline vermis. Optogenetic stimulation consisted of 100 Hz stimulation (5 ms pulse width) using a 465 nm LED (Doric Lenses). Recordings were repeated across multiple tracks over a period of no more than 4 days. Between recordings, the craniotomy was protected using silicone elastomer (Kwik-Sil, World Precision Instruments)

### Behavioral Testing

*B*ehavioral testing was conducted during the animal light cycle and in a dark behavior room. To reduce stress and low activity levels during the light cycle, mice were transferred to the behavior room at least 30 minutes prior to testing. To reduce handling stress, animals were familiarized with the behavioral arena as well as fiber optic attachment protocols for at least 2 days prior to testing (7-15 minutes per familiarization session).

Innate fear behavioral testing was performed in a 25 × 25 × 25 cm acrylic behavioral chamber with an LCD monitor positioned 40 cm above the arena floor (**Figure 1C**; as in (Carroll *et al*., 2025). Mice were familiarized with the behavioral chamber for at least 2 days (∼5-7 minutes per day) prior to innate fear testing. On test day(s), mice were placed into the chamber for at least 2-3 minutes prior to the presentation of the first looming stimulus. The looming stimulus consisted of a high contrast, rapidly expanding disk (**Figure 1D**; 0° to 20° visual angle in 333 ms) in the center of the arena (Yilmaz & Meister, 2013; Carroll *et al*., 2025). Each individual trial of the looming stimulus consisted of 5 loom repetitions across ∼6 seconds. Trials were manually triggered using a Master-8 pulse generator (AMPI), allowing for trial initiation during periods of movement. In a subset of experiments, optogenetic stimulation of vermal Purkinje cells was provided either just prior to the onset of the looming stimulus (preceding the loom by ∼500 ms) or 2-5 minutes prior to the presentation of the looming stimulus. Mice that received optogenetic stimulation were acclimated to the fiber optic attachment during familiarization trials. Unless otherwise noted, optogenetic stimulation (470 nm LED) consisted of 1 second of 100 Hz stimulation (5 ms pulse duration). Optogenetic stimulation followed numerous different protocols including coincident optogenetic stimulation during loom presentation on all trials, following only trial 1, and prior to loom presentation.

Open field locomotion was assayed in the same behavioral arena. As above, mice were first familiarized with the arena for two days, including placement of the fiber optic cannula, followed by a single day of testing. On test day, at least 5 minutes of baseline activity were recorded prior to optogenetic stimulation. Optogenetic stimulation consisted of 5 separate epochs (470 nm light, 100 Hz, 5 ms pulse duration, 1 second total duration), delivered randomly over a ∼2.5 minute window (∼30 second intervals). Following stimulation, behavior was monitored for an additional 5 minutes, allowing for within-animal comparisons of locomotion before and after vermal Purkinje cell stimulation.

To evaluate whether cerebellar stimulation resulted in aversive or appetitive behaviors, we used a real-time place preference (RTPP) assay. RTPP testing was conducted in a standard three-chambered apparatus consisting of two visually distinct (stripes vs. dots) rooms, which were connected by a clear a front-facing plexiglass hallway (**Figure 1E**). We utilized the same RTPP timeline as described by (Bimpisidis *et al*., 2020), consisting of two chamber familiarization days, two RTPP testing days (context A), followed by a conditioned recall (CR) day. Then, after a 3-day break, the cycle was repeated with RTPP pairing to the opposite context (context B), allowing for paired-context reversal testing. The RTPP experiments were counter-balanced, with half of the animals receiving stimulation within the striped room and the other half within the dotted room for “context A”. Behavioral sessions were initiated with mice being placed in the neutral hallway and access to the other two rooms blocked. After 1 minute of acclimation following handling, the room dividers were removed, and 15 minutes of behavioral testing would begin. On RTPP days, optogenetic stimulation (470 nm; 100 Hz, 5 ms pulse duration, 1 second total duration) was provided upon an animal’s initial entry into the paired contextual room and only reissued upon exit and reentry. On familiarization, pre-test, conditioned recall days, animals were still attached to the optic cable, but no instances of photo-stimulation were applied.

### Data Analysis

For all behavioral experiments, mouse behavior was recorded using an infrared camera (raspberry pi or Doric behavioral camera). To track animal position and velocity for innate fear and open-field testing we analyzed videos using DeepLabCut (DLC) marker-less pose estimation software. Following pose estimation, data was analyzed using a custom Python analysis pipeline. For each frame of the video, the x- and y-position of the mouse’s center of mass was identified. The animal speed was calculated frame-by-frame by dividing the change in animal position by the interval frame rate. Velocity data was smoothed using a rolling average across 10 frames, and immobility was defined as any 500 ms period during which the animal velocity was less than 2 cm/sec (Carroll et al., 2025). Percent time spent immobile was then calculated within the 20 s window after the onset of visual looming stimulus. To evaluate time spent in the center versus the perimeter of the chamber, we used a custom Python code that defined the center 80% of the chamber floor against the remaining 20% of the perimeter. This code used the same frame-by-frame location data from DLC providing analysis on the percent time that was spent in the center of the chamber, the distance traveled, and the velocity of an animal before and after photo-stimulation. To evaluate the percentage of time spent in each chamber during the real-time place preference assay, we used the location tracking software ezTrack (Pennington *et al*., 2019, 2021). This software compares each frame in a video against an empty chamber control frame to evaluate the animal’s location across time.

### Histological Staining

In a subset of behavioral experiments (n = 23), we confirmed cannula localization within the cerebellar vermis. Random mice from each cohort underwent a trans-cardiac perfusion to histologically evaluate fiberoptic cannula locations. Animals were anesthetized, perfused with 4% paraformaldehyde, and 80 μm coronal cerebellum slices were obtained using a Precisionary Instruments Compresstome. Cerebellar slices were mounted and stained using a cresyl violet stain to confirm anatomical location (**Figure 1B**). Cannula placement was confirmed using an Olympus APEXVIEW fluorescence microscope at 4X magnification. The location of the cannula was then confirmed using Paxinos and Franklin’s mouse brain atlas.

### Statistical testing

Data are reported as mean ± S.E.M unless otherwise noted. In all figures, symbol orientation reflects animal sex (males: upward triangles, females: downward triangles) Data analysis and statistical testing were completed in GraphPad Prism software. Statistical comparisons between two groups were calculated using either a paired or unpaired two sample t-test, as indicated in the text. For multi-trial data, a repeated measure one-way or two-way ANOVA was performed, as appropriate, followed by post-hoc comparison testing, if warranted. To quantify habituation, we subtracted the immobility on Trial 3 from the immobility on Trial 1, which was then compared to a hypothetical mean of zero (indicating no change in immobility across trials) in a one sample t-test. Unless otherwise noted, all experiments were performed with a cohort of at least 10 animals that was sex-balanced when transgenic mouse availability allowed. The n values reported reflect the number of animals in each experiment.

## Results

### Optogenetic stimulation of vermal Purkinje cells silences FN neurons

To begin to understand how acute changes in cerebellar activity may influence innate fear behavior, we first characterized, *in vivo*, how optogenetic stimulation of vermal Purkinje cells modulates spiking activity in the downstream fastigial nucleus (FN). To do this, we made loose cell-attached single unit recordings from neurons in the FN while optogenetically stimulating vermal Purkinje cells at the brain surface in awake, head-fixed mice. To confirm that our recordings were made from neurons in the cerebellar nuclei, we selected cells with high spontaneous firing rates that did not show evidence of complex spikes (Mercer *et al*., 2016; Vaaga *et al*., 2020). The basal firing rate in males and females was the same when measured *in vivo* (**Figure 2A-B**; males: 68.2±6.2 sp/s, n = 24 cells; females: 78.8±6.4 sp/s, n = 23 cells; unpaired t-test: p = 0.24; t(45) = 1.19), despite differences in spontaneous firing seen *in vitro* (Mercer *et al*., 2016; Vaaga *et al*., 2020). We next tested the extent to which optogenetic stimulation of vermal Purkinje cells modulates the firing rate of FN neurons. Cells from both males (n = 1) and females (n = 2) were pooled for firing rate modulation analysis. As expected, optogenetic stimulation (500 ms, 100 Hz, 5 ms pulse duration) significantly reduced the firing rate of FN neurons without a significant increase in post-stimulation rebound firing (**Figure 2C-D**; baseline period: 77.3±7.6 sp/s; optogenetic stimulation: 23.3±5.3 sp/s; post-stimulation period: 94.8±9.1 sp/s; RM one-way ANOVA: p < 0.0001; n = 12 cells; F(1.635, 17.99) = 48.36 ; Dunnett’s multiple comparison test: baseline vs. opto stim: adjusted p = 0.0045; baseline vs. recovery: adjusted p = 0.017). Two putative mCbN recordings were removed from analysis as optogenetic stimulation did not reduce mean firing rate, which was an a *priori* exclusion criteria for identifying mCbN units. While the degree of suppression gradually decreased throughout the duration of the stimulation, the firing remained suppressed relative to baseline firing rates throughout stimulation. Collectively, these data indicate that 100 Hz optogenetic stimulation of vermal Purkinje cells effectively and rapidly modulates spike output from the FN, resulting in a significant perturbation of cerebellar output.

**Figure 2:**
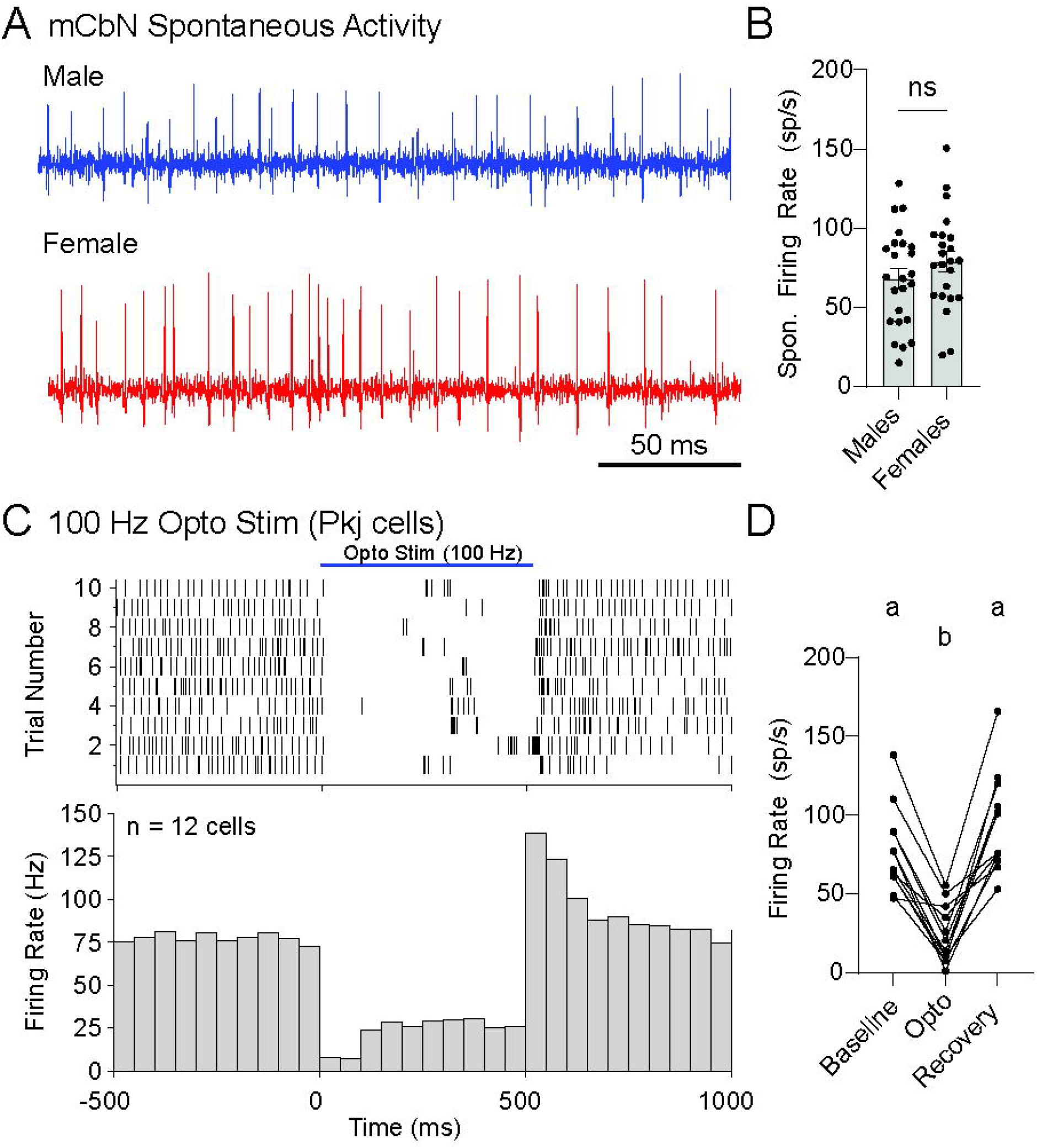
Optogenetic stimulation of Purkinje cells in the cerebellar vermis reduces firing rate of FN neurons. (**A-B**) Spontaneous firing rate of unlabeled neurons in the FN in male (A, blue) and female (A, red) L7-cre::Ai32 mice. (**B**) Distribution of the spontaneous firing rate in both the male (blue) and female (red) FN neurons. (**C**) Spike raster plot (upper) and corresponding peristimulus time histogram (lower) of FN neuron firing rate before, during, and after 100 Hz optogenetic stimulation of vermal Purkinje cells. A majority of FN neurons reduced their firing rate during optogenetic stimulation. (**D**) Line plot of firing rate changes in FN cell activity before, during, and after optogenetic stimulation of vermal Purkinje cells.

### Optogenetic disruption of cerebellar activity alters innate fear responses

The FN directly projects to the midbrain periaqueductal gray (PAG; (Gonzalo-Ruiz *et al*., 1990; Vaaga *et al*., 2020; Frontera *et al*., 2020); furthermore, we previously found that within the PAG, cerebellar input modulates the strength of both synaptic excitation and inhibition in freezing-related neurons (Vaaga *et al*., 2020). More specifically, cerebellar input increases the strength of inhibition while decreasing the strength of excitation on freezing-related premotor neurons, thereby having a net inhibitory effect on PAG circuit activity. At the behavioral level, repeated presentation of looming threats results in robust behavioral habituation (i.e. reduced freezing) across trials (Tafreshiha *et al*., 2021; Carroll *et al*., 2025). Together, these observations raise the possibility that cerebellar-induced synaptic modulation within the PAG may serve as the synaptic substrate for behavioral habituation by reducing the efficacy with which afferent synaptic input drives spike output in the PAG.

To test this prediction, we optogenetically stimulated vermal Purkinje cells to suppress FN output during presentation of a looming visual threat that elicits robust freezing behaviors (Yilmaz & Meister, 2013; De Franceschi *et al*., 2016; Tafreshiha *et al*., 2021; Carroll *et al*., 2025). First, to obtain baseline behavior while controlling for any direct responses to light, we used C57Bl/6J mice that do not express ChR2 and applied identical optogenetic stimulation. In these mice, looming stimuli elicited an increase in the percent time immobile (**Figure 3B**; baseline: 21.3±4.3%; looming: 66.2±4.7%; n = 16 mice; paired t-test < 0.0001; t(15) = 10.50), similar to that observed in non-optogenetically stimulated C57BL/6J mice (Carroll *et al*., 2025). To characterize the habituation profile, we repeated the same optogenetic stimulation + looming stimulus across three total trials separated by ∼5 minutes. As expected, in the wildtype mice, repeated presentation of the looming stimulus resulted in habituation across repeated looms (**Figure 3C**; RM one-way ANOVA: p = 0.028; F(1.496, 22.45) = 4.690).

**Figure 3:**
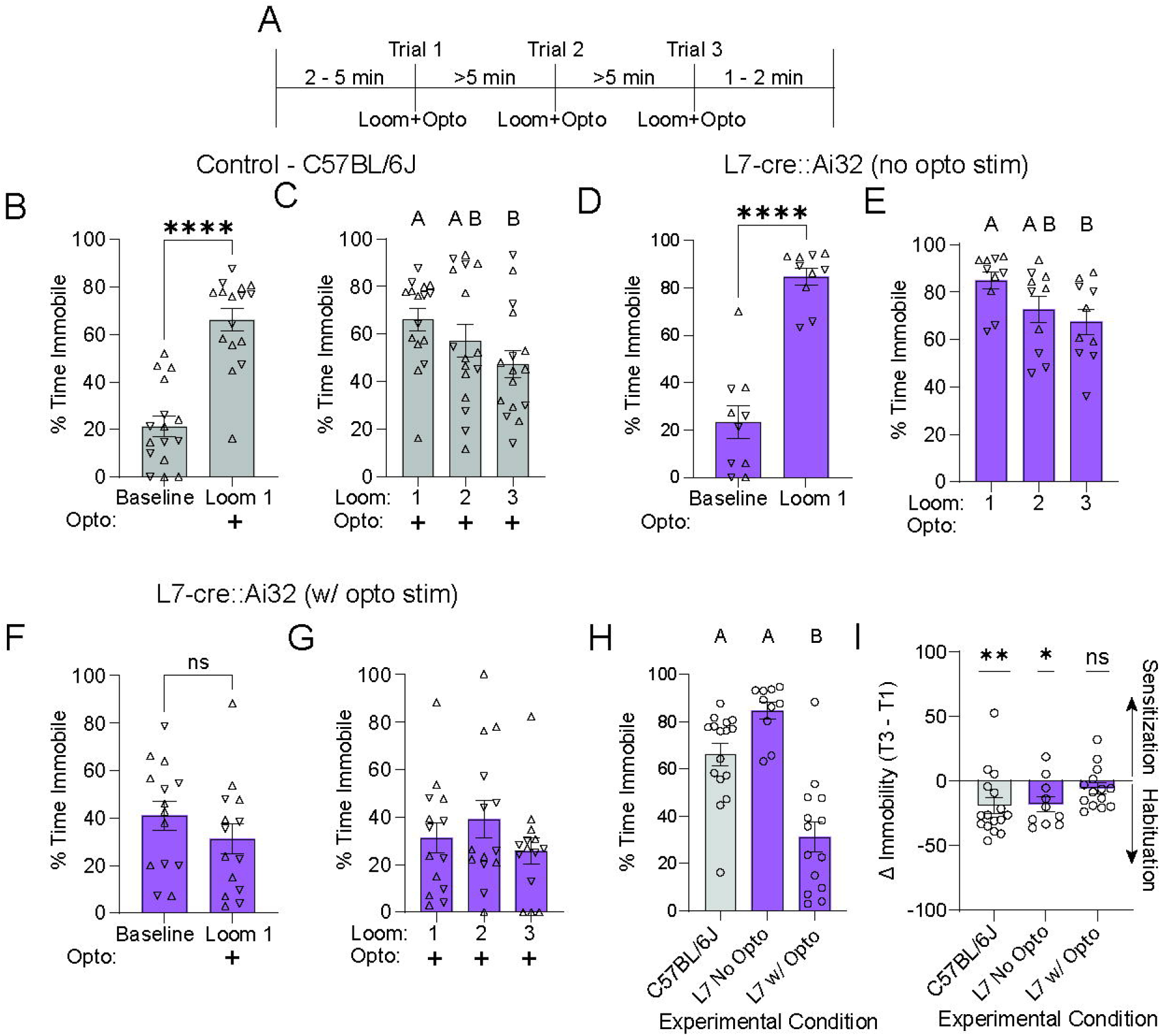
Optogenetic manipulation of cerebellar output impairs innate fear habituation. (**A**) Experimental timeline for innate fear task. Three loom trials were presented at ∼5 minute intervals, with each trial paired with optogenetic stimulation. (**B-C**) Control cohort (C57BL/6J wildtype mice) immobility responses. Looming visual stimuli elicits a significant increase in immobility on the first trial (B), which habituates across repeated trials (**C**). (**D-E**) Innate fear responses in L7-cre::Ai32 mice without optogenetic stimulation of Purkinje cells. (**F-G**) Innate fear responses in L7-cre::Ai32 mice with optogenetic stimulation. (**H**) Comparison of immobility response on Trial 1 across experimental conditions. (**I**) Comparison of the change in immobility across repeated trials, compared to a null hypothesis of 0, indicating no change in immobility. For all figure panels, sex is indicated by symbol (males: upward triangle; females: downward triangle).

We next evaluated freezing responses in our experimental mice (L7-cre::Ai32) without cannulation or optogenetic stimulation to test for potential genetic differences in fear responsivity. As in wildtype animals, looming stimulation elicited an immobility response (**Figure 3D**; baseline: 23.2±6.9%; looming: 84.7±3.6%; n = 10 mice; paired t-test < 0.0001; t(9) = 7.89). Additionally, repeated presentation resulted in the expected pattern of habituation across trials (**Figure 3E**; RM one-way ANOVA: p = 0.02; F(1.520, 13.68) = 5.654). Collectively, these data indicate similar patterns of immobility and habituation are observed under conditions with intact cerebellar signaling (e.g. wildtype mice with optogenetic stimulation and L7-cre::Ai32 without optogenetic stimulation).

To specifically test the effect of cerebellar perturbation on looming-evoked freezing, we optogenetically stimulated vermal Purkinje cells at the onset of the looming stimulus in L7-cre::Ai32 mice. Strikingly, optogenetic disruption of cerebellar output significantly reduced looming-evoked freezing responses (**Figure 3F;** baseline: 41.0±6.1%; looming: 31.1±6.4%; n = 14 mice; paired t-test = 0.26, t(13) = 1.19). We next investigated the effect of disrupting cerebellar output on innate fear habituation. Because optogenetic perturbation of cerebellar activity at the onset of each looming stimulus resulted in little freezing, there was no significant habituation observed across trials (**Figure 3G**; RM one-way ANOVA: p = 0.23; F(1.332, 17.32) = 1.616). We further directly compared immobility responses on the first loom trial across conditions, which revealed a significant overall effect of experimental condition on immobility, driven by reduced freezing in mice with disrupted cerebellar output (**Figure 3H**; one-way ANOVA: p <0.0001, F(2, 37) = 24.41). Together, these observations suggest that intact cerebellar output at the onset of the looming stimulus is required for mice to engage in looming-evoked freezing responses (Koutsikou *et al*., 2014). It is worth noting that vermal stimulation resulted in a motor response during stimulation consisting of full body extension and backwards locomotion. Despite this motor phenotype, the duration of the looming stimulus (∼6 seconds) outlasted the duration of the optogenetic stimulation (1 second), providing sufficient time for the animal to observe and respond to the looming threat. Further, in wildtype mice looming stimulation results in 35-60 second epochs of freezing, far outlasting the duration of the stimulus itself (Carroll et. al., 2025). Therefore, the motor phenotype alone is unlikely to be sufficient to account for the reduced freezing response.

To directly compare and quantify habituation across conditions, we calculated the Δ immobility by subtracting the immobility on Trial 1 from Trial 3 (i.e. T3 – T1), allowing us to test the null hypothesis that there was no change in immobility across trials (i.e. Δ immobility = 0). In both wildtype and L7-cre::Ai32 mice, the Δ immobility was significantly less than the null hypothesis of 0 (**Figure 3I**; C57BL/6J: -18.8±6.2, one-sample t-test: p = 0.0086, t(15) = 3.018; L7-cre::Ai32 (no opto): -17.6±5.9, one sample t-test: p = 0.0154, t(9) = 2.983) indicating habituation. In contrast, the Δ immobility in L7-cre::Ai32 mice with optogenetic stimulation did not significantly differ from a hypothetical mean of 0 (**Figure 3I**; L7-cre::Ai32 (w/ opto): -5.3±4.3, one sample t-test: p = 0.23, t(13) = 1.261), indicating a lack of habituation across trials. Collectively, these data suggest that disrupting cerebellar activity impairs innate fear habituation across repeated trials.

The occlusion of freezing responses in mice with disrupted cerebellar output precludes a single interpretation, as the observed lack of habituation may reflect an inability to engage in the behavior itself during optogenetic stimulation. Therefore, to begin to disambiguate the acute effects on immobility and any potential effects of cerebellar silencing on habituation, we next tested whether transient optogenetic stimulation has long lasting impacts on fear behaviors. To do this, we modified the experimental design to isolate the optogenetic manipulation to a single trial (Trial 1), leaving the remaining two trials as loom-only (**Figure 4A**). Consistent with our previous results, optogenetic stimulation during the looming stimulus significantly resulted in attenuated freezing responses (51.0±8.2% immobility). Interestingly, however, subsequent presentations of the looming stimulus alone resulted in significantly elevated immobility responses (**Figure 4B**; Trial 2: 76.6±3.1% immobility; Trial 3: 69.8±7.4% immobility; RM one-way ANOVA: p = 0.018; n = 10 mice; F(1.524, 13.71) = 6.050). Interestingly, there was no evidence of habituation between Trials 2 and 3 (**Figure 4B**; Tukey’s post-hoc comparison: p = 0.5356). Comparing the immobility on Trial 3 vs. Trial 1, the data points tended to cluster above the unity line (**Figure 4C**), indicating that the immobility on Trial 3 was greater than that observed on Trial 1. However, the Δ immobility score did not significantly differ a hypothetical value of 0 (-18.8±9.5, one-sample t-test: p = 0.077; t(9) = 1.99). Together this data suggests that disrupting cerebellar output disrupts innate freezing only on trials with concurrent stimulation, however, even transient disruptions of cerebellar output may result in a long-lasting change in innate fear.

**Figure 4:**
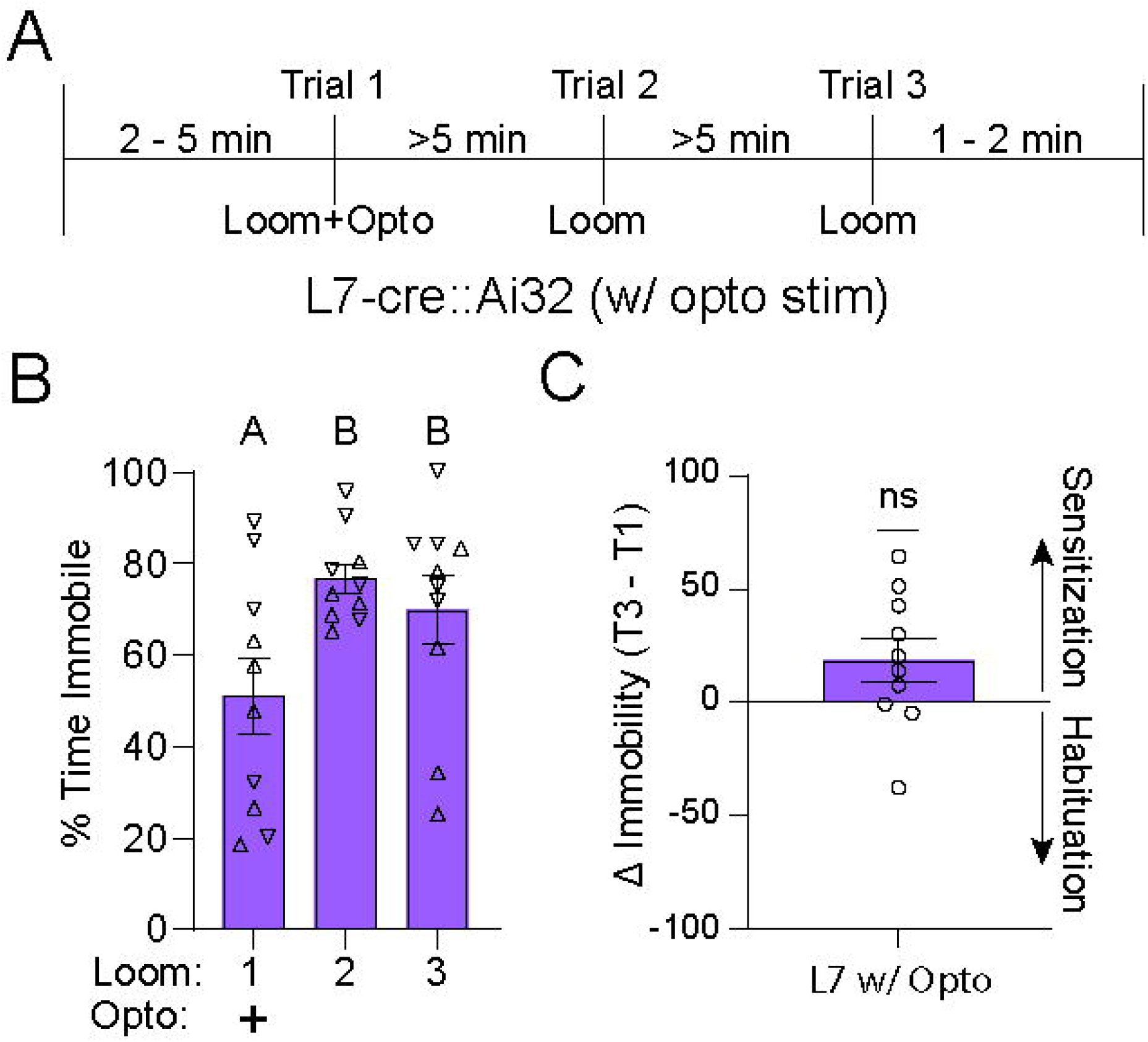
Single trial optogenetic stimulation results in a potentiated fear response. (**A**) Experimental timeline of three loom trials, with optogenetic stimulation only on trial 1. Each trial was separated by ∼5 minutes. (**B**) Experimental cohort (L7-cre::Ai32) immobility responses across repeated trials. (**C**) Change in immobility across repeated trials

### Perturbations of cerebellar activity result in long-lasting changes in fear habituation

We next sought to determine whether the apparent fear sensitization observed following cerebellar stimulation persisted over longer time periods, as the previous experiments were done with a 5-minute inter-stimulus interval. We previously demonstrated that in wildtype mice, longer inter-trial intervals (24 hours) results in an accelerated habituation phenotype characterized by a larger decrease in immobility between Trials 1 and 2, which then remains largely stable between Trials 2 and 3 (Carroll et al., 2025). We utilized a similar experimental design in which optogenetic manipulation of cerebellar circuits was restricted to Trial 1, and innate fear responses were re-tested across 3 subsequent test days (Trials 2 – 4) at 24-hour intervals (**Figure 5A**). Consistent with previous findings, wildtype mice displayed a significant decrease in the time immobile between trials 1 and 2, which was maintained at low levels of immobility across remaining trials (**Figure 5B**; RM one-way ANOVA: p = 0.0007; n = 10 mice; F(2.576, 23.19) = 8.827).

**Figure 5:**
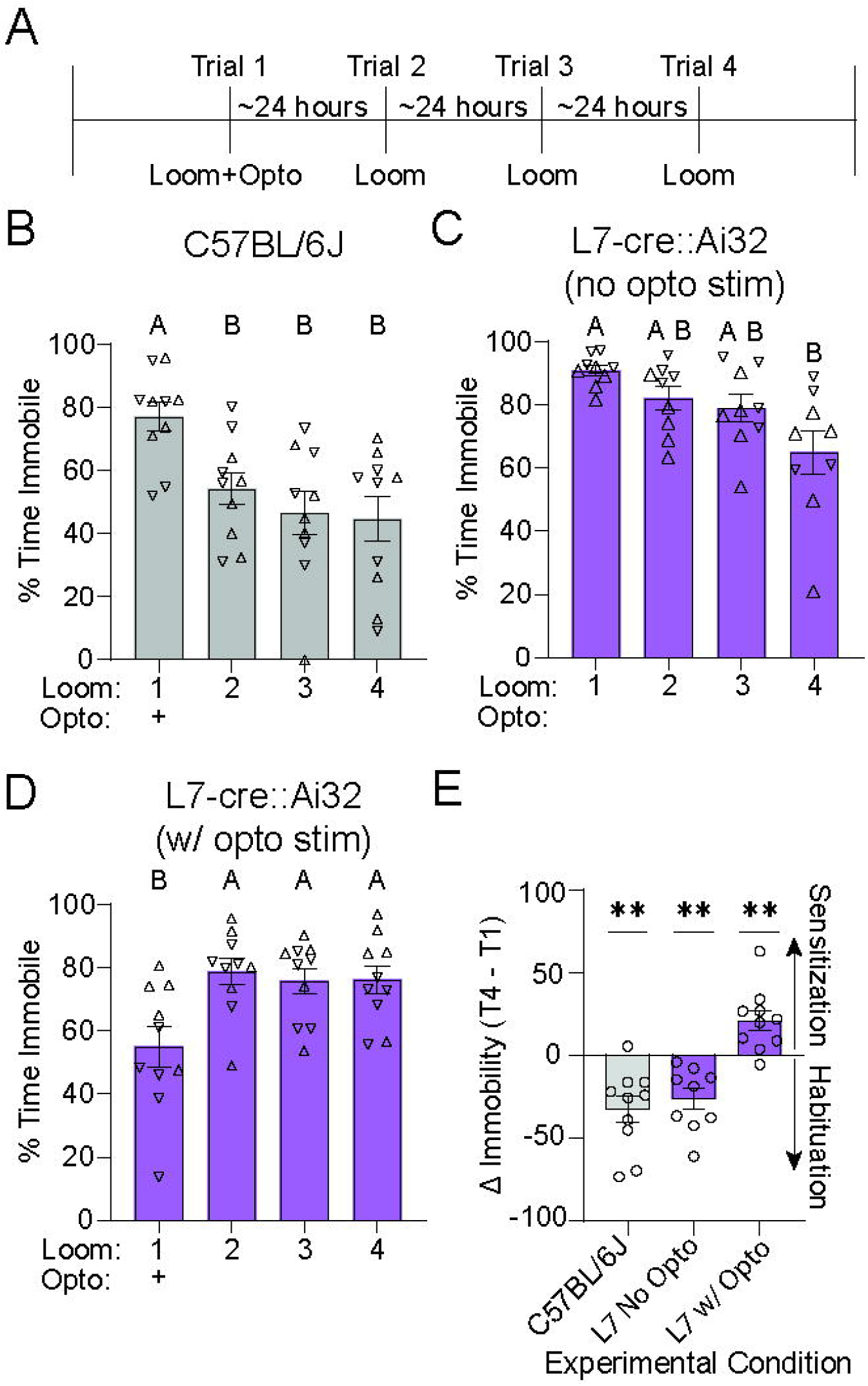
Cerebellar stimulation results in a prolonged increase in innate fear responsivity at increased time intervals. (**A**) Experimental timeline consisting of four loom trials, with optogenetic stimulation on trial 1 only. Each trial was separated by ∼24 hours. (**B-D**) Immobility responses across trials in control (C57BL6J) mice (**B**), L7-cre::Ai32 mice without optogenetic stimulation (**C**) and L7-cre::Ai32 mice with optogenetic stimulation (**D**). (**E**) Comparison of the change in immobility across trials across experimental conditions.

Consistent with this behavioral pattern, the Δ immobility metric was significantly less than 0 (**Figure 5E**; -32.4±7.8; one sample t-test: p = 0.0025, t(9)=4.14), indicative of habituation. Interestingly, the pattern of habituation differed somewhat in the L7-cre::Ai32 control cohort (without cannulation or optogenetic stimulation). More specifically, the L7-cre::Ai32 mice without optogenetic stimulation showed significant habituation across trials, but the responses habituated more gradually across successive days (**Figure 5C**; RM one-way ANOVA: p = 0.0032; n = 9 mice; F(3, 32) = 5.639). Despite this altered pattern, these mice showed significant habituation across trials (**Figure 5E**; Δ immobility: -26.1±6.4; one sample t-test: p = 0.0035, t(8)=4.091). Together, these data suggest that mice with intact cerebellar signaling engage in behavioral habituation across longer timescales, although the specific pattern of habituation may be altered in L7-cre::Ai32 transgenic mice.

In contrast, the mice with optogenetic perturbation of cerebellar output (restricted to Trial 1) showed a significant *increase* in immobility between trials 1 and 2, consistent with reduced immobility during coincident optogenetic stimulation and looming visual stimulation. Interestingly, however, subsequent trials without optogenetic stimulation resulted in stable, strong immobility responses across testing days that did not show evidence of habituation across unstimulated trials (**Figure 5D**; RM one-way ANOVA: p = 0.0015; n = 10 mice; F(1.941, 17.47) = 9.736). Consistent with an overall fear sensitization across trials, the Δ immobility was significantly greater than a hypothetical mean of 0 (**Figure 5E**; 21.3±6.0; one-sample t test: p = 0.0065, t(9) = 3.522). To explicitly measure whether responses remained stable between Trials 2 and 4, we compared the Δ immobility between Trial 4 and Trial 2 (T4-T2). The Δ immobility between these trials did not significantly differ from a hypothetical mean of 0 (-2.6±4.1; one-sample t test p = 0.54, t(9)=0.643), reinforcing the observation of stable immobility responses in trials following optogenetic perturbation. Collectively, this data suggests that even a single epoch of vermal Purkinje cell optogenetic stimulation decreases the propensity to undergo habituation on subsequent loom-only trials, an effect which is maintained for at least 72 hours.

As a final method of disambiguating the effects of cerebellar stimulation on fear habituation, we completely de-coupled the optogenetic stimulation from the presentation of the looming stimulus. To do this, mice were placed in the open field arena for a baseline period of ∼5 minutes followed by a ∼2.5 minute period which included 5 sets of optogenetic stimulation (100 Hz, 1 second duration each) delivered at random (∼30 second) intervals. Approximately 90 seconds after the final optogenetic stimulation, mice were exposed to the looming visual stimulus (**Figure 6A**). In both control (**Figure 6B**) and experimental cohorts (**Figure 6D**), looming triggered robust freezing responses, indicated by a significant increase in the percent immobility after the stimulus (control cohort: baseline: 23.5±3.7%; looming: 78.2±7.0%; n = 9 mice; paired t-test: p < 0.0001; t(8): 7.13; experimental cohort: baseline: 35.6±7.3%; looming: 79.3±4.3%; n = 10 mice; paired t-test: p = 0.0001; t(9): 6.56). Further, the initial immobility response on the first trial did not significantly differ across control and experimental cohorts (**Figure 6F**; unpaired t-test: 0.89, t(17)=0.135). The high freezing observed in the experimental cohort suggests that cerebellar stimulation *during* the looming stimulus impairs the ability of the animal to freeze; however, freezing remains intact following *isolated* disruptions of cerebellar activity.

**Figure 6:**
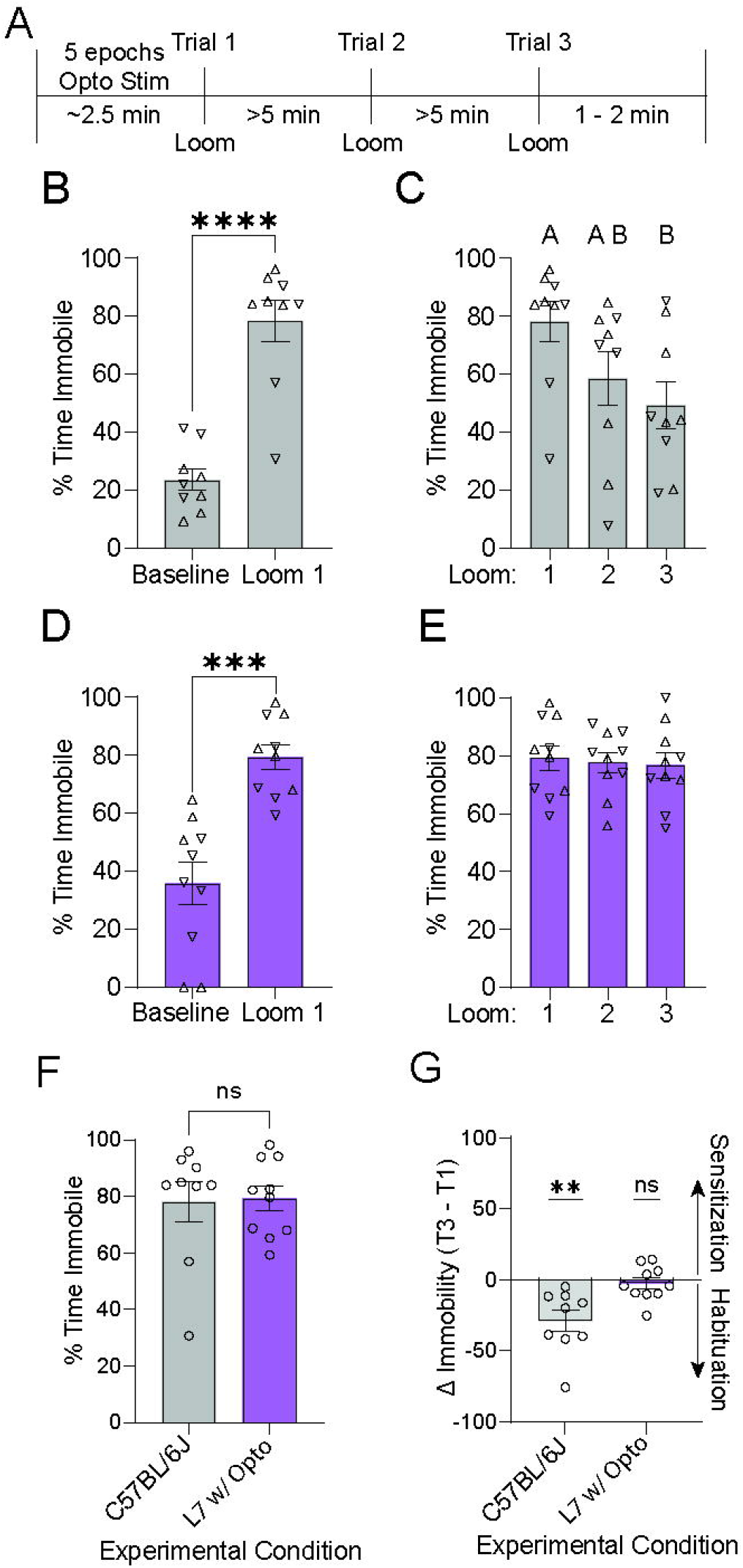
Cerebellar optogenetic stimulation alone results in an enhanced fear state. (**A**) Experimental timeline of isolated optogenetic stimulation followed by three loom trials, with each trial separated by ∼5 minutes. (**B-C**) Control cohort immobility response across repeated trials. (**D-E**) Immobility responses in L7-cre::Ai32 mice following optogenetic stimulation of vermal Purkinje cells prior to loom presentation. (**F**) Comparison of immobility responses on Trial 1 across experimental conditions. (**G**) Comparison of the change in immobility across trials in each experimental condition.

Intriguingly, the response across repeated trials significantly differed across wildtype and experimental mice. Consistent with previous datasets, mice in the control cohort showed similar levels of habituation across repeated trials (**Figure 6C**; RM one-way ANOVA: p = 0.0049; F(1.954, 15.63) = 7.683), resulting in a negative Δ immobility (**Figure 6G**; -28.9±7.4, one sample t test: p = 0.0046, t(8)=3.89). However, disruption of cerebellar activity prior to loom exposure completely eliminated habituation in the experimental cohort (**Figure 6E**; RM one-way ANOVA: p = 0.84; F(1.795, 16.16) = 0.154), resulting in a Δ immobility that did not significantly differ from a hypothetical mean of 0 (**Figure 6G**, -2.6±3.9; one sample t test p = 0.52, t(9)=0.676). The complete lack of habituation across repeated trials suggests that cerebellar stimulation results in an enhanced fear state, in which animals maintain an elevated fear response rather than engage in adaptive habituation across trials (Carroll et al., 2025).

### Cerebellar stimulation is inherently aversive

The elevated fear state observed after transient disruption of cerebellar activity suggests that vermal Purkinje cell stimulation may, in itself, be aversive. To formally test this prediction, we first examined the effect of cerebellar stimulation on overall locomotion in the open-field arena. We recorded locomotion during a 5-minute window before and after optogenetic stimulation (**Figure 7A**; 5 epochs, 100 Hz, 1 second each). Consistent with an anxiogenic phenotype, mice with cerebellar silencing showed a significant reduction in their overall locomotion in the 5-minute period following optogenetic stimulation compared to mice in the control cohort (**Figure 7B, C**; two-way RM ANOVA: main effect of genotype: P=0.0005; n = 10 mice per group; F(1, 18) = 18.16; C57BL6J mice: pre-stim: 20.6±1.3 m; post-stim: 20.3±1.2 m; Sidak multiple comparison test: adjusted p = 0.98; L7-cre::Ai32 mice: pre-stim: 19.9±1.2 m; post-stim: 13.8±1.1 m; Sidak multiple comparison test: adjusted p = 0.018). To further characterize anxiety-like behaviors we measured the percent time spent in the center 80% of the arena; a repeated measures two-way ANOVA revealed a significant genotype by session interaction (two-way RM ANOVA: genotype x session interaction: p < 0.0001). More specifically, in control mice we observed an increase in the time spent in the center of the chamber post-stimulation, likely reflecting a decrease in anxiety across the ∼15-minute session (**Figure 7D**; pre-stim: 19.3±3.04%; post-stim: 37.1±3.0%; Sidak multiple comparison test: adjusted p = 0.0002). Conversely, in the experimental cohort, mice showed a significant decrease in the time spent in the center, consistent with an anxiogenic phenotype (**Figure 7D**; pre-stim: 29.7±4.2%; post-stim: 13.8±3.4%; Sidak multiple comparison test: adjusted p = 0.0005). Finally, as a last measure of anxiety we examined the total time spent immobile and the average velocity before and after optogenetic stimulation. Both measures remained stable in the C57BL6J mice. However, in the experimental cohort, we observed a significant increase in immobility (**Figure 7E**; two-way RM ANOVA: main effect of genotype: p = 0.015; F(1, 18) = 7.493) and an associated decrease in average velocity following cerebellar stimulation (**Figure 7E**; two-way RM ANOVA: main effect of genotype: p = 0.0024; F(1, 18) = 12.39). Taken together, these results suggest that cerebellar stimulation is aversive, resulting in an increase in anxiety-like behaviors in the open field.

**Figure 7:**
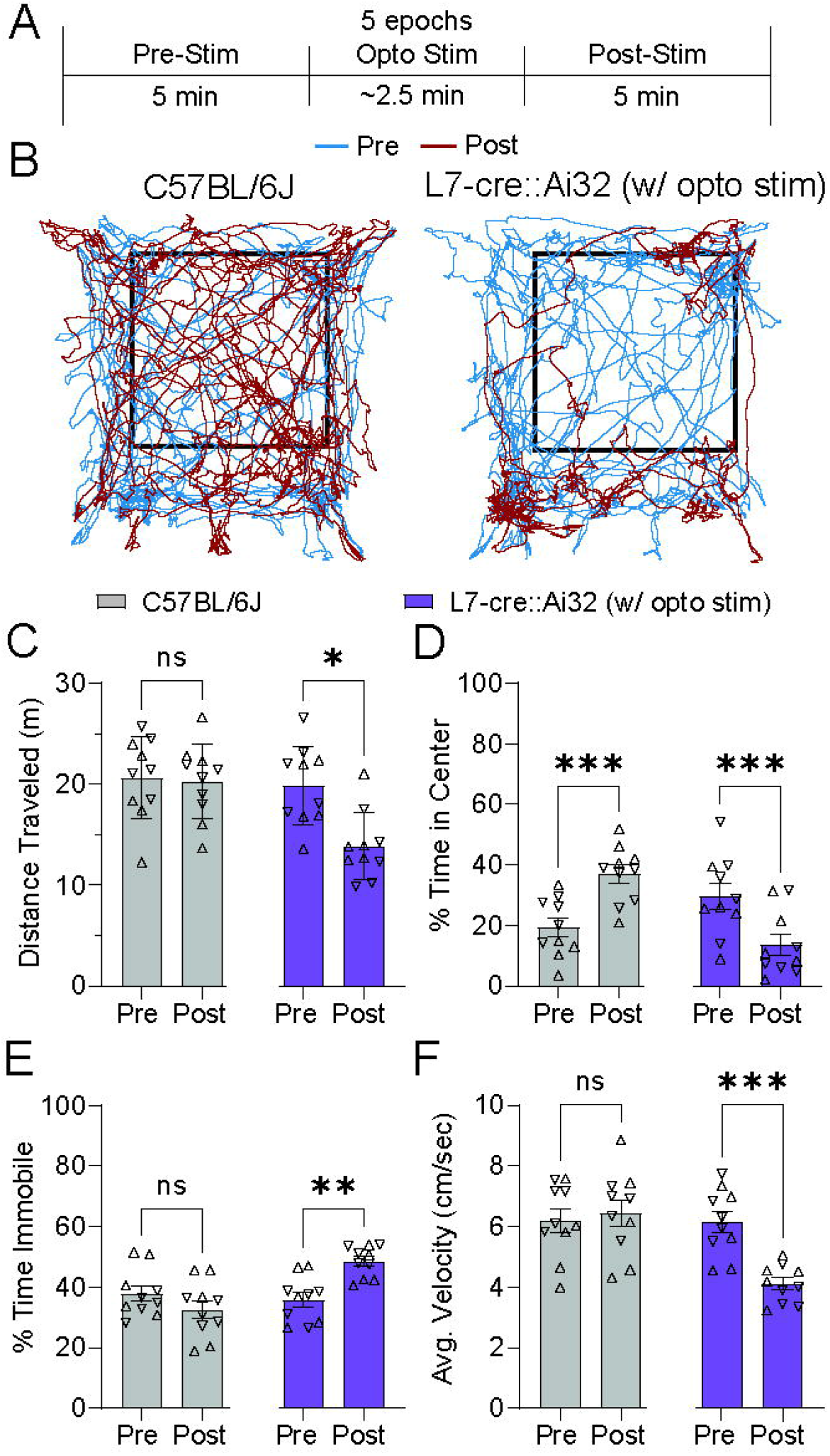
Effect of cerebellar stimulation on open field locomotion and exploration. (**A**) Experimental timeline of single-trial open field exploration before and after optogenetic stimulation. (**B**) Locomotion traces from a representative animal from control (left) and experimental (right) cohorts before (blue) and after (red) optogenetic stimulation. Black square demarcates the boundary between the center (80%) and perimeter (20%) of the chamber. (**C-F**) Evaluation of anxiety-like behaviors including distance travelled (**C**), percent time in center (**D**), percent time immobile (**E**) and average velocity (**F**) in control (grey) and experimental (purple) cohorts.

Finally, to directly test whether vermal Purkinje cell stimulation was aversive, we utilized a real-time place preference assay to directly couple cerebellar optogenetic stimulation with behavioral choice. To do this, we used a three-chamber behavioral arena with two visually distinct, large rooms and a connecting plexiglass antechamber (**Figure 1E**). All mice were exposed to a counter-balanced, blocked trial design which included chamber familiarization, optogenetic pairing with Context A, and reversal testing by optogenetic pairing with Context B (**Figure 8A**). Following each context pairing, behavior was additionally evaluated during a conditioned recall test session (CR1 and CR2) where no additional stimulation was applied. Importantly, on trial and reversal days, mice only received optogenetic stimulation (1 second, 100 Hz) upon each *entry* to the paired chamber.

**Figure 8:**
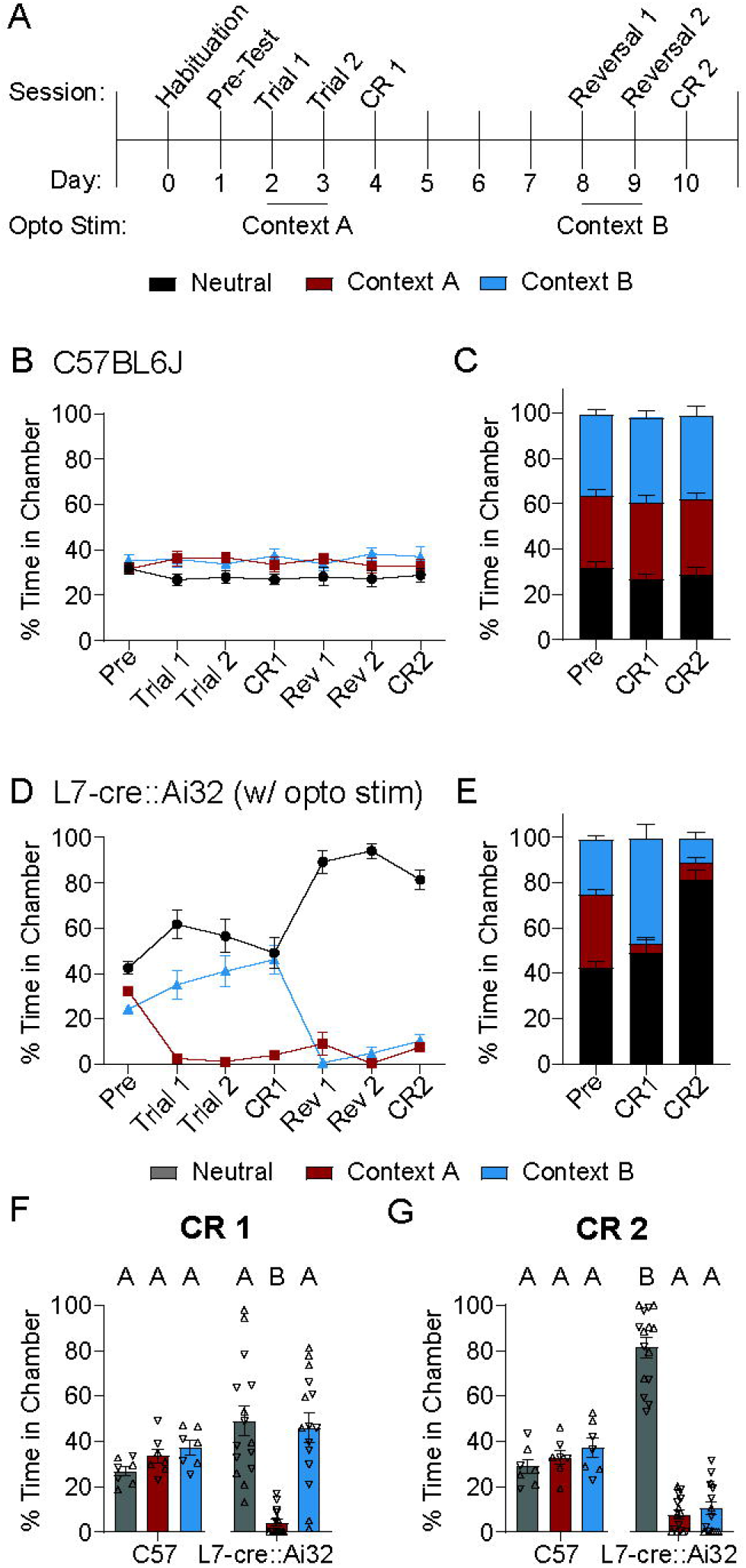
Cerebellar stimulation results in strong place aversion. (**A**) Experimental timeline of real-time place preference behavioral assay (see methods section for session descriptions). (**B, C**) Comparison of time spent in each chamber per session in control cohort. Line graph (**B**) displaying all behavioral testing days and bar graph (**C**) comparing pre-test and conditioned recall (CR) days. (**D, E**) Comparison of time spent in each chamber per session in experimental cohort. Line graph (**D**) displaying all behavioral testing days and bar graph (**E**) comparing pre-test and CR days. (**F, G**) Comparison of time spent in each chamber in control and experimental cohorts on CR days.

As expected, the control cohort showed no preference across the three chambers, spending an equal amount of time in each chamber across all test days (**Figure 8B, C**; two-way RM ANOVA: main effect of Chamber: p = 0.06; Session x Chamber interaction: p = 0.68). Conversely, in the experimental cohort, although there was no significant preference during the pre-test session, optogenetic stimulation paired with entry to Context A resulted in a rapid decrease in the percent time spent in the paired chamber as early as the first paired trial (**Figure 8D, E**; two way RM ANOVA: main effect of Chamber: p <0.0001; Session x Chamber interaction: p < 0.0001; pre-test: 32.4±1.9%; Trial 1: 2.6±1.0%). This apparent aversion was maintained across trials with reinforcement and persisted during the first conditioned recall session (CR1: 4.1±1.5%).

Perhaps more strikingly, optogenetic contingency reversal, in which the optogenetic stimulation was paired with entry to Context B, did not result in a behavioral reversal. Instead, mice began avoiding entry into Context B (pre-test: 24.3±1.7%; Reversal 1: 0.7±0.5%), while maintaining their aversion to Context A (Reversal 1: 9.1±5.2%). This resulted in mice spending a significantly increased amount of time in the neutral antechamber (89.3±5.1%) rather than entering either context which had been previously paired with optogenetic stimulation. To quantify these changes, we examined the time spent in each chamber during sessions without optogenetic reinforcement (i.e. conditioned recall (CR) 1 and 2). During both CR1 and CR2, mice in the control cohort spent equal time in each chamber. Conversely, during CR1, mice in the experimental cohort showed a significant reduction in the time spent in context A (**Figure 8F**; CR1 neutral chamber: 49.1±6.8; paired chamber: 4.1±1.5%; unpaired chamber: 46.2±6.4%; Tukey’s multiple comparison test: neutral vs. paired chamber: p<0.0001; neutral vs. unpaired chamber: p = 0.93). Further, during CR2, mice in the experimental cohort showed a significant reduction in the time spent in either Context A or Context B (**Figure 8G**; CR2 neutral chamber: 81.4±4.3%; initially paired chamber: 7.6±2.0%; reversal paired chamber: 10.3±2.9%; Tukey’s multiple comparison test: neutral vs. initially paired chamber: p < 0.0001; neutral vs. reversal paired chamber: p < 0.0001). These results strongly suggest that optogenetic stimulation of the cerebellar vermis results in strong real-time place avoidance, which is maintained across numerous days and resistant to reversal.

## Discussion

Collectively, our results suggest that optogenetic activation of vermal Purkinje cells disrupts adaptive innate fear behaviors through two apparent mechanisms. First, optogenetic stimulation of vermal Purkinje cells during looming stimuli disrupts the ability of mice to engage in normal freezing responses and attenuates habituation across repeated trials. Second, optogenetic stimulation prior to looming stimulation results in an enhanced fear state, in which mice maintain abnormally high levels of freezing across repeated trials. Consistent with this observation, our results suggest that independent of any effects on innate fear processing, stimulation of the cerebellar vermis is anxiogenic and aversive – resulting in reduced locomotion in the open field and driving robust real-time place preference aversion.

### Limitations of the present study

It is worth noting a few limitations of our study. First, optogenetic activation of Purkinje cells produces a robust motor response during stimulation, directly opposing engagement in freezing behaviors which are, by definition, the lack of movement. It is important to note, however, that the visual looming stimulus lasts for ∼6 seconds whereas the optogenetic stimulation only occurred for 1 second. It is therefore highly likely that following optogenetic stimulation, mice would have sufficient time to observe and respond to the looming stimulus and engage in (delayed) freezing. Our results instead suggest that following brief optogenetic stimulation, mice largely ignore the looming stimulus, suggesting that normal cerebellar activity may be required at the *onset* of the looming stimulus in order to engage in appropriate fear responses (Koutsikou *et al*., 2015). The reduced freezing during optogenetic stimulation also limited our ability to interpret disruptions in trial-to-trial habituation, which we circumvented using alternative study designs to limit trials with cerebellar stimulation or de-couple optogenetic stimulation from looming stimuli altogether.

Further, it is worth noting that the fear responses in the L7-cre::Ai32 mouse line without optogenetic stimulation were slightly different from wildtype controls. While the L7-cre::Ai32 mice demonstrated robust initial fear responses and normal habituation on short time intervals, the experimental mice showed reduced habituation at 24-hour intervals. This reduction in habituation at long time scales may reflect persistent changes in neural activity within the cerebellum in the L7-cre::Ai32 mouse line, especially if long-term cerebellar dependent plasticity is a central component to long-term fear learning (Sacchetti *et al*., 2002, 2004; Strata *et al*., 2011; Frontera *et al*., 2020; Lawrenson *et al*., 2022). Despite this potential confound, the L7-cre::Ai32 mice did show significant habituation across trials, in contrast to conditions with optogenetic stimulation in which habituation was not observed.

Finally, our optogenetic manipulation was targeted to the midline vermis rather than regions of the vermis known to receive input from the PAG (Watson *et al*., 2013; Koutsikou *et al*., 2014). However, our goal in this set of experiments was to manipulate cerebellar output by changing activity in the fastigial nucleus, not to specifically disrupt afferent PAG-related activity in the cerebellar cortex. Therefore, the decision to stimulate Purkinje cells through midline vermal stimulation, in theory, allows for intact PAG-driven activity in the cerebellar cortex while disrupting cerebellar output more broadly within the nuclei.

### Cerebellar interactions with downstream limbic circuits

The cerebellum has complex interactions with multiple nodes in the distributed limbic system (Apps & Strata, 2015), including mono or di-synaptic connectivity with the periaqueductal gray, amygdala, and prefrontal cortex (Snider & Maiti, 1976; Gonzalo-Ruiz *et al*., 1990; Middleton & Strick, 1997, 2001; Vaaga *et al*., 2020; Frontera *et al*., 2020). Perhaps the most direct interaction with the fear network comes via monosynaptic connectivity with multiple cell types within the ventrolateral midbrain periaqueductal gray (Vaaga *et al*., 2020; Frontera *et al*., 2020). Of note, the apparent predominant functional impact of cerebellar stimulation in the vlPAG is mediated by activation of a local population of dopaminergic neurons (Vaaga *et al*., 2020). Dopaminergic activation favors synaptic inhibition onto freezing premotor neurons – strengthening IPSCs while attenuating EPSCs. While these synaptic interactions may favor habituation through reduced synaptic activation of freezing-related premotor neurons, this view is complicated by emerging *in vivo* work, suggesting that the vlPAG plays more complex and nuanced roles in fear processing – including computations of threat probability and the generation of a positive prediction error (Wright & McDannald, 2019; Walker *et al*., 2019; Wright *et al*., 2019; Strickland & McDannald, 2022). These observations raise the possibility that cerebellar input to the vlPAG modulates these processes, which may account for altered patterns of habituation across repeated trials by altering fear appraisal mediated by vlPAG circuits. Further, while there is strong evidence that the PAG is downstream of the cerebellum, there is additional evidence that freezing behaviors require PAG activation of the cerebellum itself (Watson *et al*., 2013; Koutsikou *et al*., 2014). This suggests that, like many other circuits, the cerebellum forms a multi-synaptic loop with the PAG. Such loops may facilitate both direct roles of the cerebellum in driving motor responses and participating in cerebellum dependent learning to update behavioral responses under conditions in which there is a mismatch between perceived threat and actual danger.

Our results support this general framework. Disrupting normal cerebellar output during looming stimuli disrupted the ability of mice to engage in appropriate freezing responses. Such disruption may reflect a direct role of the cerebellum in driving freezing motor responses. Alternatively, cerebellar activation may be required for the activation of freezing-related premotor neurons in the vlPAG, a subset of which do receive direct cerebellar input (Vaaga *et al*., 2020). Further, uncoupling optogenetic stimulation from looming stimuli resulted in an enhanced fear state with little to no habituation across repeated trials. These findings suggest a potential role for cerebellar circuits in fear appraisal and/or computations related to threat probability.

It is also possible that the observed effects of cerebellar stimulation are not entirely mediated by connectivity between the cerebellum and the PAG. Stimulation of the cerebellum also results in short latency responses in the amygdala (Snider & Maiti, 1976), which is a well-known hub for regulation of both conditioned and innate fear responses (Herry *et al*., 2008; Ciocchi *et al*., 2010; Tovote *et al*., 2016; Fadok *et al*., 2017). Recent evidence suggests that cerebellar circuits are di-synaptically coupled to the basolateral amygdala, providing a circuit architecture by which activity in the cerebellum may modulate amygdala circuitry (Jung *et al*., 2022). Additionally, the cerebellum and prefrontal cortex are reciprocally connected (Brodal, 1978; Hartmann-von Monakow *et al*., 1981; Leichnetz *et al*., 1984; Middleton & Strick, 1997, 2001; Schmahmann & Pandya, 1997; Kelly & Strick, 2003) providing another pathway by which cerebellar activity may regulate innate fear. While the specific role of the prefrontal cortex in innate fear has not been well-established, the prefrontal cortex provides dense innervation of the periaqueductal gray, suggesting a role in regulating fear responses (An *et al*., 1998; Vertes, 2004; Gabbott *et al*., 2005; Skog *et al*., 2024; Lukinic *et al*., 2025). Collectively, these observations strongly support a role for the cerebellum in modulating limbic function, but complicate straight-forward, mechanistic interpretations of cerebellar action.

### Cerebellar contributions to non-motor functions in health and disease

Some of the early evidence that the cerebellum broadly contributes to non-motor function comes from clinical literature – in which damage to the cerebellum can result in a constellation of affective and cognitive symptoms referred to as the Cerebellar Cognitive Affective Syndrome (CCAS; (Schmahmann, 2004, 2021; Hoche *et al*., 2018; Ahmadian *et al*., 2019). Of note, damage to the cerebellar vermis is most often associated with emotional dysregulation including flat affect and a general blunting of emotional responsivity (Supple *et al*., 1988; Schmahmann & Sherman, 1998). These clinical observations suggest that the cerebellum, and more specifically the vermis/fastigial (medial) cerebellar nucleus contributes to emotional processing, which may include fear responses. Additionally, the cerebellum has been implicated in a variety of psychiatric illnesses and fear disorders, including phobia, anxiety and panic disorders, and PTSD, where notable differences in cerebellar volume and resting state connectivity are observed compared to healthy controls (De Bellis & Kuchibhatla, 2006; Baldaçara *et al*., 2011; Lanius *et al*., 2017; Holmes *et al*., 2018; Reid, 2022).

While the role of the cerebellum in non-motor functions in animal studies has become increasingly recognized in recent years, the specific contribution of cerebellar circuits to emotional regulation remains elusive. One central question which has emerged is the extent to which there exists a universal cerebellar transform (Schmahmann, 1991; Schmahmann & Caplan, 2006; Schmahmann *et al*., 2019), in which cerebellar contributions to sensorimotor transformations can be applied to non-motor computations. The near crystalline microanatomy of the cerebellum suggests that the cerebellar cortex performs qualitatively similar computations across task domains (Voogd & Glickstein, 1998; Apps & Garwicz, 2005). In this view, cerebellar computations during innate fear behaviors may be conceptualized as cerebellar-driven learning during threat-danger mismatch. Simply put, if the perceived threat does not match reality (i.e. under conditions in which there is no real danger; a threat-danger mismatch), it may trigger a climbing-fiber mediated error signal which serves to update threat appraisal; such computations would be similar to climbing fiber mediated error signals updating a motor program. Such computations may then be shared with downstream circuitry, including the PAG, in accordance with its role in computations of threat probability and generation of threat-related positive prediction errors (Wright & McDannald, 2019; Walker *et al*., 2019). Our results are broadly consistent with this framework. Specifically, we demonstrate that the cerebellum contributes to both fear expression and more broadly to adaptive habituation across repeated trials. However, the bidirectional effect of cerebellar perturbation on these processes (namely disrupting freezing expression and enhancing fear state) suggest that the cerebellum may play a key role in computing potential a mismatch between perceived and actual danger. This is consistent with a broader framework in which cerebellar circuits contribute to prediction error-based updating of threat appraisal, extending canonical cerebellar learning principles into the domain of emotional behavior.

## Acknowledgements

We are grateful to Indira Raman for her mentorship and partial grant support for the initial experiments done by CEV in the Raman lab (R35-NS116854).

